# An intranasal OMV-based vaccine induces high mucosal and systemic protecting immunity against a SARS-CoV-2 infection

**DOI:** 10.1101/2021.08.25.457644

**Authors:** Peter A. van der Ley, Afshin Zariri, Elly van Riet, Dinja Oosterhoff, Corine P. Kruiswijk

## Abstract

The development of more effective, accessible and easy to administer COVID-19 vaccines next to the currently marketed mRNA, viral vector and whole inactivated virus vaccines, is essential to curtain the SARS-CoV-2 pandemic. A major concern is reduced vaccine-induced immune protection to emerging variants, and therefore booster vaccinations to broaden and strengthen the immune response might be required. Currently, all registered COVID-19 vaccines and the majority of COVID-19 vaccines in development are intramuscularly administered, targeting the induction of systemic immunity. Intranasal vaccines have the capacity to induce local mucosal immunity as well, thereby targeting the primary route of viral entry of SARS-CoV-2 with the potential of blocking transmission. Furthermore, intranasal vaccines offer greater practicality in terms of cost and ease of administration. Currently, only eight out of 112 vaccines in clinical development are administered intranasally. We developed an intranasal COVID-19 subunit vaccine, based on a recombinant, six proline stabilized, D614G spike protein (mC-Spike) of SARS-CoV-2 linked via the LPS-binding peptide sequence mCramp (mC) to Outer Membrane Vesicles (OMVs) from *Neisseria meningitidis*. The spike protein was produced in CHO cells and after linking to the OMVs, the OMV-mC-Spike vaccine was administered to mice and Syrian hamsters via intranasal or intramuscular prime-boost vaccinations. In all animals that received OMV-mC-Spike, serum neutralizing antibodies were induced upon vaccination. Importantly, high levels of spike-binding immunoglobulin G (IgG) and A (IgA) antibodies in the nose and lungs were only detected in intranasally vaccinated animals, whereas intramuscular vaccination only induced an IgG response in the serum. Two weeks after their second vaccination hamsters challenged with SARS-CoV-2 were protected from weight loss and viral replication in the lungs compared to the control groups vaccinated with OMV or spike alone. Histopathology showed no lesions in lungs seven days after challenge in OMV-mC-Spike vaccinated hamsters, whereas the control groups did show pathological lesions in the lung. The OMV-mC-Spike candidate vaccine data are very promising and support further development of this novel non-replicating, needle-free, subunit vaccine concept for clinical testing.

## INTRODUCTION

Since the outbreak of SARS-CoV-2^1^, development of a vaccine against this emerging pathogen has been given the highest priority globally. The first vaccine, a mRNA vaccine^2^, entered the market with unprecedented speed early December 2020^3^. Soon other mRNA, viral vector and whole inactivated vaccines followed. All COVID-19 vaccines on the market today and the majority of COVID-19 vaccines in development are intramuscularly administered, targeting the induction of systemic immunity. Although the current vaccines protect against emerging SARS-CoV-2 variants, efficacy wanes^4^. Therefore, the development of additional vaccines aims to broaden and strengthen the immune response because of concerns about reduced vaccine-induced immune protection to emerging variants^5,6^, the fact that many people in large parts of the world do not have access to vaccines yet, and there is still limited knowledge on the longevity of current vaccine responses.

Intranasal vaccines have the capacity to induce not only systemic but also local mucosal immunity. In view of the respiratory tropism of the SARS-CoV-2 virus, a vaccine delivered intranasally, able to elicit mucosal immunity directly at the port of entry of the virus and preventing transmission, would be desirable^7-11^. Mucosal protection is the first line of defense in the human body against respiratory pathogens, and intranasal antigen delivery is associated with induction of mucosal immunity including production of IgA antibodies and the priming of T cells in nasal tissues. Furthermore, intranasal administration is a needle-free vaccination route which could aid in mass vaccination campaigns by increasing the ease and speed, by decreasing costs and reducing pain associated with vaccination. Currently only eight out of 112 vaccines in clinical development are intranasally administered and only two of those are subunit vaccines^12^. The other intranasal vaccines in clinical development are mainly vector vaccines for which there is a risk that pre-existing immunity against the vector can impair the magnitude of the elicited immune response after vaccination or booster vaccination. The vaccine concept described here is an intranasal subunit vaccine containing recombinant spike protein and based on outer membrane vesicle (OMV) platform technology.

OMVs can bud off spontaneously from the cell surface of many Gram-negative bacteria and are non-infectious vesicles. They present surface antigens in a form that is efficiently recognized by the immune system and activate innate immunity through the presence of multiple PAMPS like lipopolysaccharide (LPS) and lipoproteins. Because of their strong immunogenicity, OMVs have been explored extensively as vaccine candidates, especially in the case of *Neisseria meningitidis*. We have optimized the meningococcal OMV platform to tailor it for safe and effective use in humans. By combining genetically detoxified *lpxL1* LPS with deletion of the *rmpM* gene to increase blebbing, high yield production and isolation of native OMV with reduced endotoxicity became possible. In multiple clinical and non-clinical toxicity and immunogenicity studies OMVs were successfully tested and proven safe and effective^13-19^. Meningococcal OMVs can also be applied as a carrier or adjuvant for antigens from other pathogens. Previously we have demonstrated how expression of the OspA lipoprotein from *Borrelia burgdorferi* on meningococcal OMV could be used to induce protection against borreliosis in a mouse model^20^. However, this application requires efficient expression of heterologous antigens in OMV, which is not always feasible as it requires compatibility with the bacterial outer membrane biogenesis machinery. As an alternative, recombinant antigens can be produced separately, and externally linked to the OMVs, enabling the use of OMV as a carrier and adjuvant for the e.g. the SARS-CoV-2 spike protein.

The SARS-CoV-2 spike protein is found in prefusion conformation on infectious virions and in this form the protein displays most of the neutralizing epitopes that can be targeted by antibodies^21^ to prevent the entry process mediated by the receptor binding domain (RBD) of the spike protein^22,23^. The ACE2 receptor on the host cell recognizes the RBD and mediates the structural transition from prefusion to postfusion conformation of the protein^24^. McLellan and coworkers rapidly developed a prefusion stabilized spike protein (S-2P), by introducing a double-proline substitution in the S2 domain of the spike protein^25^ to aid vaccine design. Several of the first vaccines developed, like the mRNA vaccines mRNA-1273^2^ and BNT162b2^26^ use the S-2P in their vaccine designs. An improved prefusion stabilized spike protein, HexaPro^27^, was developed combining four additional proline substitutions into S-2P. Hexapro showed increased stability, higher expression in mammalian cells, proper conformation, thus preserving its antigenicity and a third of the particles were found to be in a two-RBD-up conformation^27^. Hexapro is thus an ideal target to be included in the design of SARS-CoV-2 vaccines.

In the present study, we developed an innovative subunit vaccine for COVID-19 in which a HexaPro spike protein containing the D614G mutation^28-30^ is associated to OMV, that serve as a carrier and adjuvant for the spike molecule, using the short amphipathic peptide sequence mCRAMP fused to the C-terminus. Both mCRAMP as well as its human orthologue LL-37 are short amphipathic peptides that belong to the cathelicidins, vertebrate host defense peptides defined by their conserved pro-region^31^, and have a strong binding affinity for LPS. The interaction with LPS is through negative phosphate groups in lipid A and the inner core oligosaccharide, and the positively charged region of LL-37 or mCRAMP, which is followed by membrane insertion^32^. We have used this LPS-binding property to direct a recombinant Spike-mCRAMP fusion protein to associate with OMV (Figure 1).

**Figure 1.**
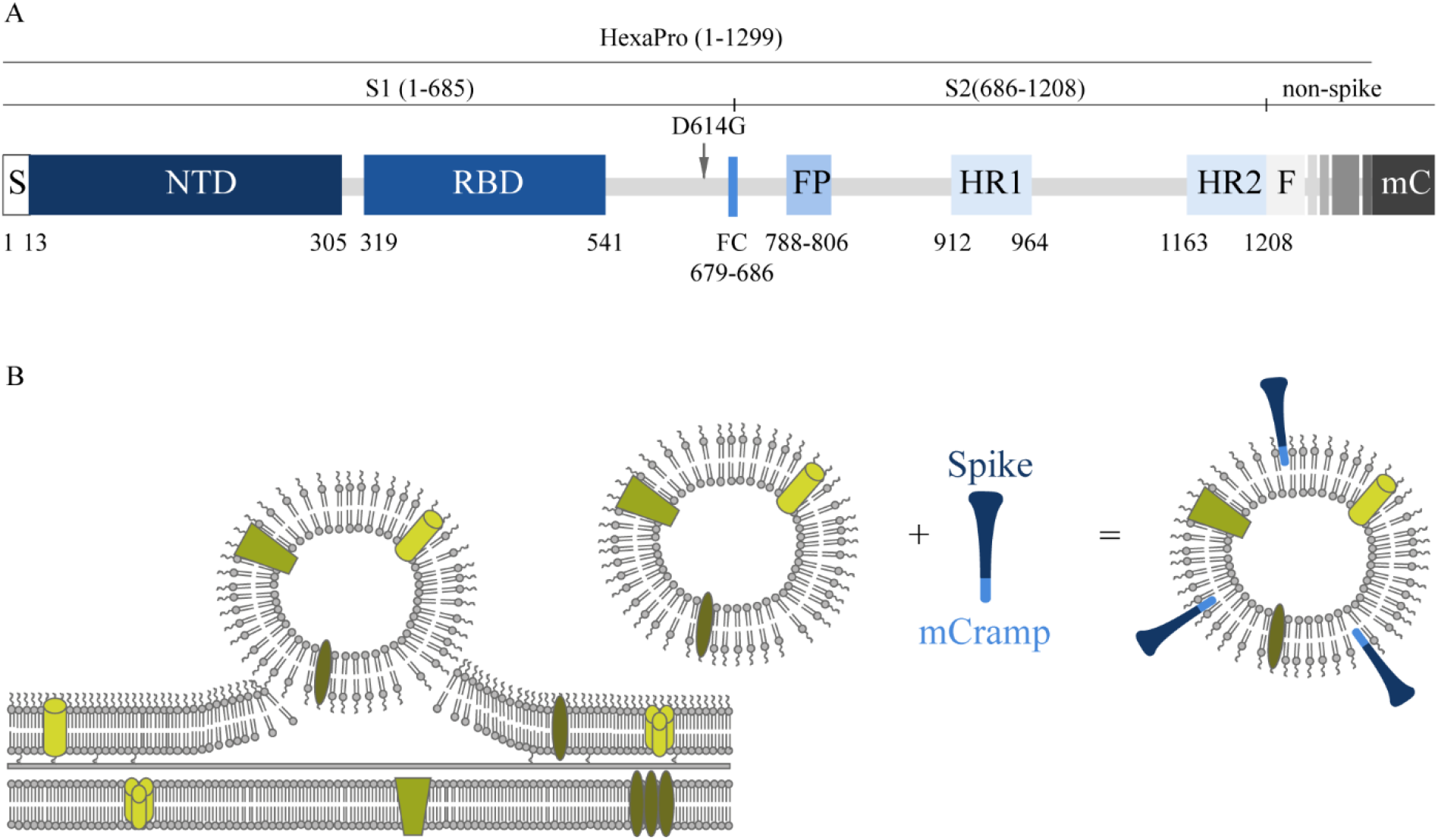
Schematic presentation of the spike molecule and OMV decoration with spike. (A) Schematic of 2019-nCoV S primary structure colored by domain. S, signal sequence; NTD: N-terminal domain; RBD: Receptor Binding Domain; FP: fusion peptide; HR1-HR2: heptad repeat 1 and 2; FC: disrupted S1/S2 furin cleavage site (R682G, R684S, R685S). Features that were added to the ectodomain (amino acid 1-1208) expression construct are colored white gray gradient (amino acid 1209-1333). From light to dark gray: Foldon trimerization motif (F); HRV 3C site; 8xHis tag; Twin Strep tag; 3x GGGS repeat; mCramp (mC). Not shown: Six stabilizing prolines in S2 at position F817P, A892P, A899P, A942P, K986P, V987P^27^. The aspartic acid at position 614 in the original HexaPro^27^ spike protein was replaced with a glycine (D614G). (B) schematic overview of how antigens can associate to the OMVs. In the spike protein an mCRAMP (antimicrobial peptide) motif was included which is depicted in light blue. This peptide associates spontaneously to the LPS of the OMV thereby attaching the spike protein (dark blue) to the OMV.

The immunogenicity of the OMV linked HexaPro spike vaccine concept was tested in a mouse model after administration via the intranasal route and intramuscular route. Although the mouse model is not the preferred model to study efficacy, as mice are not susceptible for infection with SARS-CoV-2, this model provides a good read-out on immunogenicity of COVID-19 vaccines. Subsequently for measuring protection, a Syrian hamster model was used in which animals were challenged with wildtype SARS-CoV-2 after vaccination and again both the intranasal and intramuscular routes were compared. In this hamster model we show protection of throat and lungs against respiratory COVID infection by our OMV-based non-replicating SARS-CoV-2 vaccine.

## MATERIALS AND METHODS

### Recombinant spike protein production

The prefusion stabilized HexaPro spike protein^27^ with and without a 3xGGGS spacer and mCRAMP sequence added to the C-terminus were produced by Celonic AG (Basel, Switserland) by transient expression in Expi-CHO cell cultures (ThermoFisher Scientific) as previously described^33^. In addition to the 3xGGGS spacer and mCRAMP sequence, the aspartic acid at position 614 in the original HexaPro spike protein was replaced with a glycine (D614G). The amino acid sequences of the spike proteins used are shown in Figure S1. The SARS-CoV-2 spike proteins with and without the 3xGGGS spacer and mCRAMP sequence added to the C-terminus, were named respectively mC-Spike and Spike, and were purified using Strep-tag affinity chromatography (Strep-tactinXT 4flow -IBA) according to manufacturer’s procedure. The purified protein was analyzed by native SDS-PAGE to verify its native configuration. Purity was verified by determining residual host cell protein, host cell DNA and endotoxin content.

### Meningococcal carrier strain construction

Starting material for strain construction was *N. meningitidis* HI5, a spontaneous *porA* deletion mutant of strain H44/76^34^. A spontaneous mutant resistant to streptomycin was isolated so that the strain became amenable for the use of a marker-free mutagenesis technique^35^. Strep^R^ mutants were selected by plating H44/76 Δ*porA* cells on GC medium containing 500 µg/ml Strep. Sequencing of their *rpsL* gene showed that they contained amino acid change K88R compared to the starting material, a mutation known to confer streptomycin resistance in other bacteria. The marker-free method is based on two-step selection/counterselection of a cassette that confers erythromycin resistance (Ery^R^) and streptomycin sensitivity (Strep^S^). For the first step, a 1^st^ crossover construct was designed consisting of the Ery^R^-Strep^S^ cassette, flanked by DNA bordering the region to be replaced in the *N. meningitidis* genome. A DUS sequence is added to ensure uptake of the DNA. For the counterselection step, a 2^nd^ crossover construct was designed identical to the 1^st^ crossover construct, except that the Ery^R^-Strep^S^ cassette is deleted. Positive transformants that carry the Ery^R^-Strep^S^ cassette in their genome were subjected to transformation with linearized plasmid carrying the 2^nd^ crossover construct. Selection for streptomycin resistance then resulted in the isolation of cells that have lost their genomic Ery^R^-Strep^S^ cassette as a result of recombination between linearized plasmid and the *N. meningitidis* genome. Using this technique, three deletions were introduced. The *siaD* gene was deleted, which results in loss of capsule and increased safety for laboratory handling. Subsequently, the *lpxL1* gene was deleted, resulting in penta-acylated lipid A and less reactogenic LPS. Last, the *rmpM* gene was deleted, which results in increased blebbing and higher OMV yields. After construction was finished, the final strain (renamed as Nm RLΔpA) was analyzed for the presence of the correct mutations by sequencing.

Linearized plasmids containing the 1^st^ crossover constructs were transformed into *N. meningitidis* H44/76 (Δ*porA*, Strep^R^, with previously introduced mutations) by adding ∼1 µg plasmid to a cell suspension of OD590 nm = 0.2 in vTSB (Fluka) with 0.1M MgCl_2_. Successful transformants were selected on GC-agar (BBL) supplemented with IsoVitaleX (BBL) and 5 µg/ml Ery. For the second step (deletion of the Ery^R^-Strep^S^ cassette), linearized plasmid containing the 2^nd^ crossover construct was added, and bacteria were plated on GC + 500 µg/ml Strep to select Strep^R^ clones no longer containing the Ery^R^-Strep^S^ cassette as a result of double crossover between the genome and the 2^nd^ crossover construct. Colonies with the correct antibiotic resistance phenotype (Ery^S^ and Strep^R^) were then further verified by PCR.

### OMV production

*N. meningitidis* OMVs were produced using our downscaled platform production process^36^. Bacteria were precultured from cryovials in 155 ml chemically defined medium in a 500-ml shake flask at 35°C and 200 rpm. At exponential phase of the cells at OD590 nm of ∼1.8, the whole preculture was transferred to a 5L single-use bioreactor (Eppendorf BioBlu3f) containing a total of 3.2L medium volume. The culture conditions were controlled (Eppendorf BioFlo320) and monitored at 35°C, pH7.2 by addition of 1M NaOH and 1M HCl. The dissolved oxygen tension was controlled at 30% by a headspace total flow aeration of 1L/min air and by increasing the stirrer speed (300-1000rpm) followed by replacing air by pure oxygen. Both oxygen and carbon dioxide percentages were analyzed in the off gas. The culture was harvested after ∼16 hours of cultivation, and two hours after the maximum Carbon Dioxide Evolution Rate (CER) resulting in an OD590 nm of 9. The culture in the bioreactor was cooled to 20°C before starting the concentration with a 790cm2 mPES hollow fiber with a 0.45µm pore size using the KMPi TFF system (Repligen). The concentrated biomass was diafiltrated with two Diafiltration Volumes (2DV) with 0.1 M Tris-HCl, pH 8.6. Afterwards, 10 mM EDTA in 0.1M Tris pH 8.6 was added to the biomass to start the extraction for 30 minutes at ambient temperature^37^. After extraction, EDTA was removed by diafiltration using the same TFF hollow fiber module with 0.1 M Tris, pH 8.6. TFF steps were performed at a constant flux of 20±5 LMH^36^. The extracted OMVs were separated from the biomass by high-speed centrifugation (30 min, 23,500xg). The supernatant containing the crude OMVs was transferred to a bottle and treated overnight at ambient temperature with 100U DNAse (Benzonase) and 2 mM MgCl_2_ to digest residual DNA. The OMVs were clarified by an Opticap SHC 0.5/0.2 µm pore size Poly Ether Sulfone (PES) depth filter to remove potential aggregates. Group separation was performed by Size Exclusion Chromatography on a XK50 Sepharose 4 FF column (Cytiva) and controlled by an Akta GE system. The eluent was 10 mM Tris/3% sucrose, pH 7.4. The OMV fraction was sterile filtered using a Supor EKV dead-end filter with 0.2 µm pore size and stored at 2-8°C.

### OMV characterization

The vesicle size distribution was measured with dynamic light scattering (DLS). Protein composition was analyzed by mass spectrometry. Total protein concentration was measured with the Peterson assay. The LPS content was quantified by a modified gas chromatography method, with LPS quantification based on the peak area of C14:3OH and using C12:2OH as internal standard^38^. PorB identity was confirmed by ELISA with monoclonal antibody MN15A14H6 which is specific for serotype 15.

### Vaccinations

#### BALB/c mice

(OlaHSD; Envigo, female, 8-9 weeks old at day 0) were immunized on day 0 and 21 via the intranasal or intramuscular route. On day 0, 21 and 35, blood was collected for assessment of the induction of antibodies against spike and SARS-CoV-2 specific neutralizing antibodies. On day 35 nasal washes and the lungs were also collected for IgA antibody determination. Groups consisted of 10 mice each. For intranasal immunization, a 20 µl inoculum was divided over both nostrils using a pipet. For intramuscular immunization, a 50 µl inoculum was injected into the thigh muscle. The OMV dose used was 15µg protein per immunization. The Spike and Spike mCRAMP dose used was also 15µg protein per immunization.

#### Syrian hamsters

(*Mesocricetus auratus*, RjHan:AURA (outbred), Janvier, France, SPF, male, 9 weeks old) were immunized on day 0 and 21 via the intranasal (50 µl) or intramuscular (100 µl) route at Viroclinics B.V. (Rotterdam, the Netherlands). For intranasal immunization, a 50 µl inoculum was divided over both nostrils using a pipet. For intramuscular immunization, a 100 µl inoculum was injected into the thigh muscle. The OMV dose used was 15 µg protein per immunization. The mC-Spike dose used was also 15 µg protein per immunization. During the study, animals were weighed, and blood was collected for assessment of spike specific IgG antibodies and induction of SARS-CoV-2 specific neutralizing antibodies. Three weeks after the second immunization (day 42), all animals were challenged intranasally with 10^4.0 TCID50 SARS-CoV-2, strain BetaCoV/Munich/BavPat1/2020. On day 4 post challenge half of the animals per group were euthanized by exsanguination under isoflurane anesthesia and necropsy was performed, with the remaining half of the animals following on day 7 post challenge.

### Enzyme-linked Immunosorbent Assay (ELISA)

Subclass specific antibodies in mice and hamsters were measured with an ELISA. Spike without the mCRAMP motif was used to coat high binding 96-wells flat bottom plates (Microlon) at a final concentration of 2µg/ml in PBS and incubated overnight at room temperature. After washing three times with 0.05% Tween-80 in water, plates were blocked using 0.5% BSA in PBS for 30 minutes at 37°C. Plates were washed three times with 0.05% Tween-80 in water and sera, nasal washes or lung homogenates were added to the plates and incubated for one hour at 37°C. Serial dilutions of pooled sera from each group were tested to determine the final serum dilution tested for all the sera. Serum, nasal wash, and lung homogenate dilutions used are described in the figure legends. After washing another three times the HRP conjugated antibody (Goat-anti-mouse HRP IgG 1:8000, IgG1 1:4000, IgG2a 1:4000, IgG2b 1:4000, IgG3 1:4000 and IgA 1:4000, southern Biotech and Goat polyclonal Ab anti-Syrian hamster HRP IgG conjugate 1:8000 from Abcam) was added to the plates and incubated for one hour at 37°C. TMB substrate (KPL SeraCare) was added to the plates after washing three times and after 10 minutes the reaction was stopped using 0.1 M H_2_SO_4_. The absorbance was measured on a microplate reader at 450 nm.

### Virus Neutralization Test (VNT)

The VNT was performed by a specialized clinical diagnostics service laboratory (Viroclinics B.V., Rotterdam, The Netherlands) on multiple samples collected during the preclinical studies. In short, samples are heat inactivated for 30 minutes at 56°C. Subsequently, serial two-fold dilutions of the samples are made in triplicate in 96-wells plates starting with a dilution of 1:5. The sample dilutions are then incubated with a fixed amount of virus (200 TCID50/well) for 1 hour at 37 °C leading to a starting dilution of the serum in the assay of 1:10. Next, the virus-antibody mixtures are transferred to plates with Vero E6 cell culture monolayers, followed by an incubation period of 5-6 days at 37°C. Subsequently, plates are scored using the vitality marker WST8.

### Pathology

At the time of necropsy gross pathology was performed at Viroclinics B.V. (Rotterdam, the Netherlands). All lung lobes were inspected, the percentage affected lung tissue estimated from the dorsal side, a gross pathological diagnosis described, and the left lung lobe inflated with and preserved in 10% formalin. Trachea and nasal turbinates were macroscopically evaluated and sampled for virology and histopathology. Relative lung weight was calculated. Histopathological analysis from selected tissues was performed for all animals. After fixation with 10% formalin, sections from left lung and left nasal turbinate, and gastrointestinal tract tissue were embedded in paraffin and the tissue sections were stained for histological examination. Lung tissue was analyzed and scored for presence and severity of alveolitis, alveolar damage, alveolar edema, alveolar hemorrhage, type II pneumocyte hyperplasia, bronchitis, bronchiolitis, peribronchial and perivascular cuffing. Scores are presented as sum of Lower Respiratory Tract (LRT) disease parameters.

### Viral load determination

Quadruplicate 10-fold serial dilutions were used to determine the virus titers in confluent layers of Vero E6 cells. To this end, serial dilutions of the samples (throat swabs and tissue homogenates) were made and incubated on Vero E6 monolayers for 1 hour at 37 °C. The monolayers were washed and incubated for 5 or 6 days at 37°C, and scored for CPE using the vitality marker WST8. Viral load analysis was outsourced to a specialized clinical diagnostic service laboratory (Viroclinics B.V., Rotterdam, Netherlands)

## RESULTS

### Production and characterization of HexaPro Spike-mCRAMP (mC-Spike)

The purified HexaPro Spike-mCRAMP protein was analyzed by non-reducing SDS-PAGE (results not shown). A band between 120-200 kDa was seen, as expected for the monomeric spike protein. Some additional bands of low intensity were observed in the higher molecular weight area, possibly indicating oligomers or aggregates. On native SDS-PAGE, three bands were visible, representing different oligomeric states of the trimeric spike protein. Determination of residual host cell protein showed a content of <140 ppm.

### OMV production and characterization

The carrier OMVs were prepared from a derivative of *N. meningitidis* strain H44/76 in which the immunodominant PorA protein had been removed, and LPS was detoxified by deletion of the *lpxL1* gene^39^. A deletion of *rmpM* was introduced to increase blebbing^38^, and of *siaD* to remove the capsule. The main characteristics of the produced OMVs are summarized in Table 1. PorB was by far the most abundant protein (70.9 %), and as expected PorA was completely absent. The vesicle size as determined by dynamic light scattering was around 100 nm.

**Table 1.** Characterization of OMVs.

| **Protein accession number** | **Protein Description** | **Relative abundance\*** |
| --- | --- | --- |
| **E6MZM0** | Major outer membrane protein P.IB | 70.9 wt% |
| **E6MXF3** | Outer membrane OpcA family protein | 5.4 wt% |
| **E6MW75** | Ribonuclease E | 2.3 wt% |
| **Q9ZHF3** | Type IV pilus biogenesis and competence protein PilQ | 1.1 wt% |
| **E6MUY7** | Outer membrane protein assembly factor BamA | 0.9 wt% |
| **E6MY46** | Lipoprotein | 0.6 wt% |
| **E6MYS2** | Peptidylprolyl isomerase | 0.6 wt% |
| **E6MWK7** | Curli production assembly/transport component CsgG family protein | 0.5 wt% |
| **E6MZU3** | 30S ribosomal protein S2 | 0.5 wt% |
| **E6N0K3** | IgA-specific serine endopeptidase | 0.5 wt% |

|  | Units | Value | Standard Deviation |
| --- | --- | --- | --- |
| Protein content | µg/mL | 606 | 32 |
| LPS content | µg/mL | 90 | 5.5 |
| DLS Z-average | nm | 100.3 | 1.8 |
| DLS PDI** |  | 0.110 |  |
| Zeta potential | mV | -35.4 | 1.9 |

\*Shown are the 10 most abundant proteins
\*\*DLS: dynamic light scattering; PDI: poly dispersity index

### Mouse immunogenicity study

Vaccination with OMVs combined with Spike protein was compared for constructs with and without the mCRAMP tag, named OMV-mC-Spike and OMV+Spike respectively. Empty OMVs were included as controls. Mice were immunized by either the intranasal or intramuscular route, and the experimental setup and timelines of the experiment are shown in Figure 2A. High serum IgG titers against purified spike protein were obtained with both OMV-mC-Spike and OMV+Spike, and not with OMVs alone (Figure 2B and Figure S2). Interestingly, IgG antibody responses were significantly higher after intranasal versus intramuscular immunization (Figure 2B and Figure S2). IgA antibody responses in both serum, nasal washes and lung were clearly present after intranasal immunization, but barely detectable in the intramuscular groups (Figure 2C and Figure S3).

**Figure 2.**
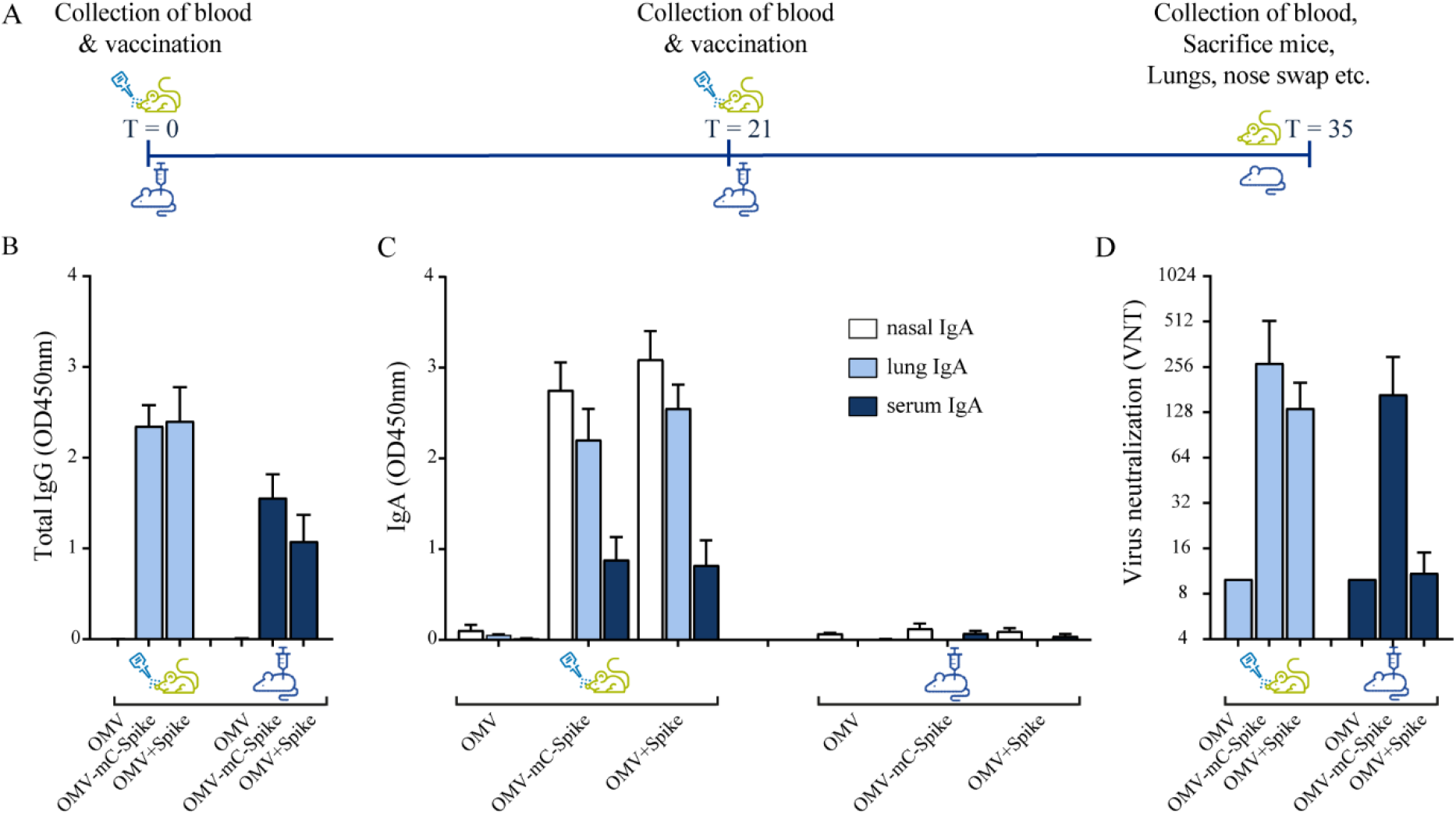
Mouse immunogenicity study. Balb/C mice were immunized intranasally or intramuscular on day 0 and day 21 with 15µg OMV (control group) or 15 µg OMV combined with 15µg Spike with the presence of mCRAMP or without mCRAMP (OMV+Spike). Sera were collected from all animals at day 35. (A) Experimental setup and timeline of the mouse immunogenicity experiment. (B) Total IgG antibody levels were measured in sera diluted at 1:50000. (C) IgA levels were measured in nasal washes (1:1 dilution), lung (1:50 dilution) and serum (1:200 dilution). (D) Virus neutralization titers were determined in sera. Data are depicted as mean ± SD and are representative results of two independent experiments.

Virus neutralization titers were only detected in the groups receiving OMVs combined with either Spike or Spike-mCRAMP protein, with the latter group (OMV-mC-Spike) showing the highest titers and highest number of responders. The difference between OMV-mC-Spike and OMV+Spike was significant for the intramuscularly but not for the intranasally immunized mice (Figure 2D and Figure S2). Statistical significance between groups is depicted in Figures S2 and S3.

### Hamster challenge study

As mice are not susceptible to SARS-CoV-2 infection, we next studied protection against viral challenge in the hamster model. Syrian hamsters were immunized with either purified mC-Spike protein or OMV-mC-Spike, with Tris-sucrose or empty OMVs as controls. The experimental setup and timelines of the experiment are shown in Figure 3A. High IgG antibody responses against spike were seen after immunization with OMV-mC-Spike for both the intranasal and intramuscular routes (Figure 3B). This was reflected in the virus neutralization titers which were also only detected in these same groups (Figure 3C). After challenge with SARS-CoV-2, almost no lung lesions were detected in the OMV-mC-Spike groups, while mC-Spike protein alone also provide significant protection compared to OMV or Tris-sucrose (Figure 3D). OMVs alone did not provide any protection, showing that generalized innate immune activation by the OMVs did not play a role. The viral load determined in throat swabs, lungs and nasal turbinates showed the highest reduction in the OMV-mC-Spike group (Figure 3E and F).

**Figure 3.**
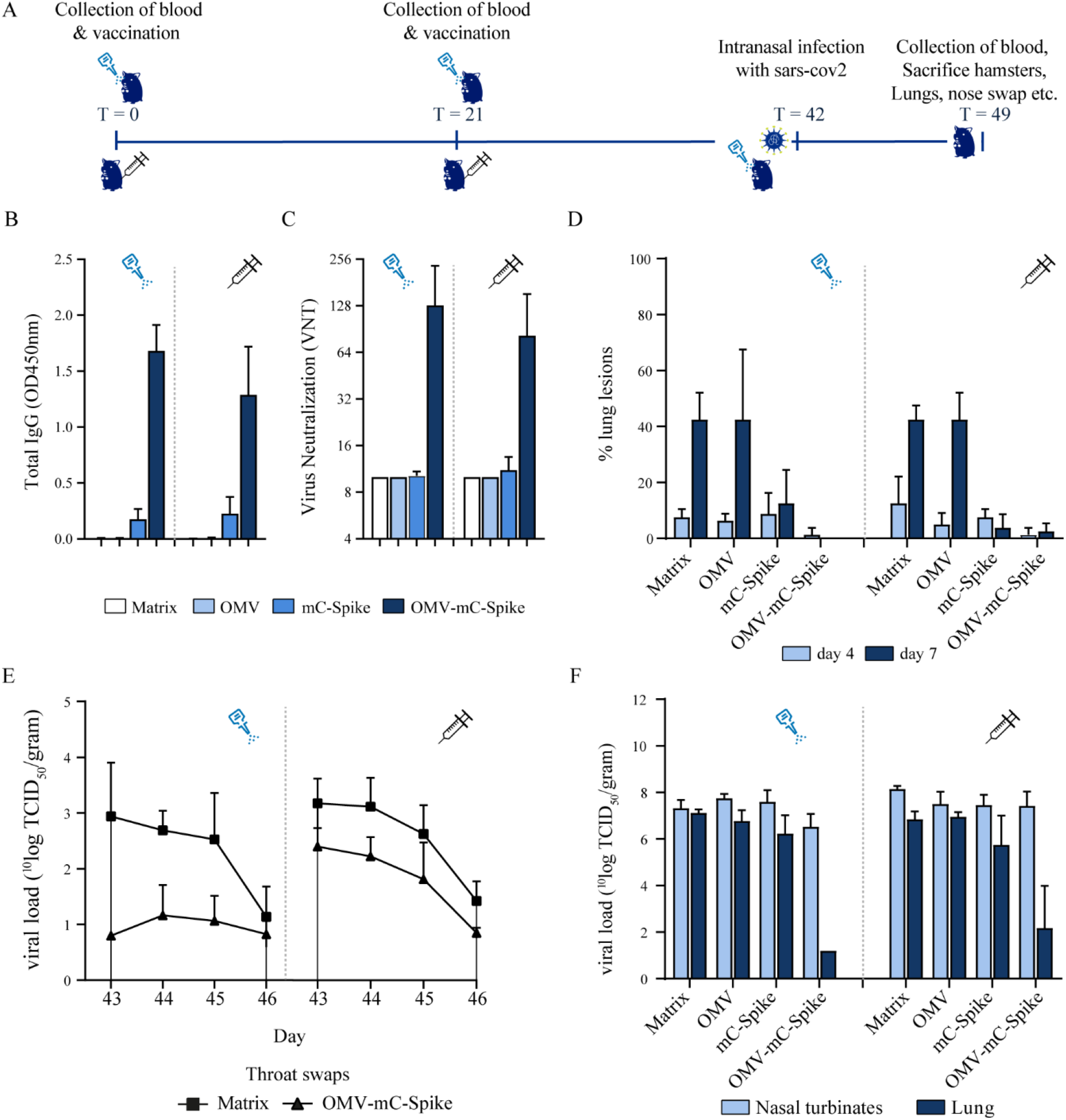
Hamster challenge study. Animals were immunized intranasally or intramuscularly on day 0 and day 21 with 15 µg OMV, 15 µg spike or 15 µg OMV and 15 µg Spike combined with mCRAMP. In the control group animals were immunized with 10mM Tris-3% sucrose, which is the OMV buffer. Sera were collected from all hamsters at experimental day 0, 21, 42, 46 and 49. At day 42, all hamsters were challenged intranasally with 10^4.0 TCID50 SARS-CoV-2, strain BetaCoV/Munich/BavPat1/2020. At day 46 half of the animals per group (4 out of 8) were sacrificed and at day 49 the remaining 4 animals were sacrificed. (A) Experimental setup and timeline of the hamster challenge experiment. (B) Total IgG anti-spike antibody levels were measured in sera (dilution 1:4000) from day 42 in an ELISA. (C) Virus neutralization was determined in sera from day 42. (D) When animals were sacrificed at day 46 (day 4 post challenge) and day 49 (day 7 post challenge) the percentage of the lung that presented lung lesions was quantified. The viral load was determined in throat swabs (E), lungs and nasal turbinates (F). Data are depicted as mean ± SD.

Vaccination was safe as the weight of the hamsters till the day of challenge was comparable for all groups (Figure S5A and S5B). After challenge the weight reduction was significantly lower for the OMV-mC-Spike group as compared to the Tris-sucrose control and the Spike protein-immunized hamsters (Figure S5C and S5D). The sum of lower respiratory tract (LRT) disease parameters as determined by gross pathology was highly reduced after intramuscular vaccination with OMV-mC-Spike, whereas after intranasal vaccination with OMV-mC-Spike no disease parameters were scored (Figure S5E). Overall, these data showed clear protection after both intranasal and intramuscular immunization with OMV-mC-Spike. Statistical significance between groups is depicted in Figure S4 and S5.

## DISCUSSION AND CONCLUSIONS

Here we describe the assessment of safety, immunogenicity and protective capacity of OMV-mC-Spike, a COVID-19 vaccine consisting of HexaPro spike that is associated with OMVs from *N. meningitidis* via a short amphipathic peptide sequence, mCRAMP, in mice and Syrian hamsters. Both intranasal and intramuscular delivery of OMV-mC-Spike induced a strong humoral response against the SARS-CoV-2 spike antigen in a 2 dose regimen. Throughout the study no adverse events were recorded in the animals, supporting the safety of the vaccine. After challenge of the hamsters with SARS-CoV-2 a clear reduction in viral loads and no lung disease burden was observed, supporting the protective capacity of OMV-mC-Spike.

All COVID-19 vaccines currently in use are administered intramuscularly, which mainly induces a systemic antibody response. However, mucosal administration of vaccines has several advantages. In general, benefits of needle-free administration include easier administration, absence of sharp waste, and convenience for the vaccinee. In addition, mucosal vaccination provides a rational reason to induce a protective immune response against pathogens, including SARS-CoV-2, that infect humans via mucosal surfaces^40^. Moreover, with respect to the immune response, different types of immune cells are activated in the mucosa, leading to a different type of both innate and adaptive immune response with a major role for IgA antibodies in the latter. IgA antibodies have several useful characteristics. For example, IgA antibodies can neutralize pathogens even before they pass the mucosal barrier, because the secretory form is transported across the epithelium. Moreover secretory IgA antibodies are polymeric, usually a dimer, which is more effective in neutralizing pathogens^40,41^, which might reduce carriage in the nose. For example, intranasal but not intramuscular vaccination of ChAdOx1 nCoV-19 vaccine reduced shedding of SARS-CoV-2 in rhesus macaques^42^. Furthermore, in contrast to IgG antibodies, IgA antibodies do not activate the complement pathway thereby limiting the induction of inflammatory responses and with respect to influenza IgA antibodies are known to be cross-protective across different strains^43^.

For SARS-CoV-2 the role of IgA antibodies and mucosal responses in general have been less intensively investigated. However, development of a mucosal vaccine is also thought to be a promising approach in the case of SARS-CoV-2^41^, since vaccination at the entry point can induce a strong local as well as systemic immune response^44,45^. Similarly to earlier findings with influenza viruses, among the induced antibodies in serum and saliva of COVID-19 patients, IgA antibodies contributed more to virus neutralization than IgG^46^. Intranasal, but not intramuscular, vaccination with OMV-mC-Spike indeed induced a high local IgA antibody response in the nose and in the lungs. Both intranasal and intramuscular vaccination with OMV-mC-Spike clearly reduced viral replication in the lungs after challenge with SARS-CoV-2. Several mucosal vaccines for humans on the market or in late stage clinical trials use live-attenuated viruses, including an oral polio vaccine (OPV), an intranasal influenza (FluMist) and a COVID-19 vaccine (COVI-VAC). As *N. meningitidis* is adapted to live in the human nasopharynx, it can efficiently interact with host cells in this site through its cell surface components, and *N. meningitidis* derived OMVs will mimic these steps making them particularly suitable for mucosal vaccination. As our OMV-based vaccine induced both local and systemic IgA antibody responses after intranasal but not intramuscular vaccination in mice, it is possible that it will not only protect against disease, but also against infection with COVID-19 thereby preventing transmission and the occurrence of variants. Intranasal vaccines can also give rise to resident memory T and B cells, thus providing additional protection against local viral replication and transmission^41^.

The results clearly demonstrate the feasibility of using the OMV platform for development of intranasal vaccines for infectious diseases and outbreaks by externally linking antigens to the OMV. Advantages of this platform are first of all the scalability of OMV production, as bacteria replicate rapidly, and the strains were genetically modified to produce a high amount of OMV. Due to the genetic detoxification of the LPS mild extraction processes of the OMV are feasible. The developed process also retains lipoproteins on the OMVs to further stimulate the immune response. Using standard filtration steps that are scalable the OMVs can be manufactured to high purity. An efficient large-scale bioreactor production process including downstream purification has been established^47^. Furthermore, OMVs are extremely stable at 2-8 °C and even elevated temperatures, and can be stockpiled, so in case of an outbreak of a pathogen, large batches of OMV can be quickly used for linking of antigens of the pathogen. Also, the platform is highly versatile, as different types of antigens, like proteins, peptides or sugars can be linked to OMVs.

In both the mouse and hamster model, virus-neutralizing antibodies are induced when the spike protein is combined with OMVs. In the hamster model, almost no lung lesions are found after challenge when vaccination was done with spike protein combined with OMVs. Adding a C-terminal mCRAMP tag to associate the spike protein with the OMVs increases the protective response in both models compared to mixing spike with OMVs. Overall, these data show that (i) Neisseria OMVs are an effective adjuvant/delivery system for the COVID-19 Spike protein, and (ii) increasing OMV association by an mCRAMP tag improves the protective response.

When using OMVs as delivery system for heterologous antigens, an important consideration is whether the antigen needs to be incorporated in the vesicles, or if just mixing is sufficient. A priori one could argue that association is advantageous, as antigens and OMVs will be taken up and processed by antigen-presenting cells simultaneously, thereby preventing induction of premature differentiation by OMVs before sufficient antigen is taken up. A study with tumor antigens and E.coli-derived OMVs showed that co-delivery was indeed more effective in inducing anti-tumor immunity in a mouse model^48^. Our previous results with the Borrelia OspA protein showed that both methods gave significant protection against Borrelia challenge in mice^49^. However, OspA is a lipoprotein and may therefore spontaneously associate to some extent with OMVs through its N-terminal lipidation anchor. Our present results with SARS-CoV-2 spike protein show that noncovalent OMV association through the mCRAMP tag is advantageous for the resulting immune response.

OMVs closely mimic the bacterial pathogens from which they are derived, and have been shown to confer a bactericidal immune response against *N. meningitidis*. Although mainly used by the intramuscular route, the size of OMVs is in the range thought to be most effective to cross the mucus^40^ and meningococcal OMVs have also been shown to induce a bactericidal response after intranasal application^50,51^. Salmonella-derived OMVs carrying the pneumococcal antigen PspA^52^ induced strong Th17-mediated immunity in mice after intranasal vaccination, leading to reduced pneumococcal colonization^53^. Intranasal immunization of mice with *Bordetella pertussis* OMVs induced mucosal IgA and Th17-mediated responses which prevented colonization of the nasal cavity and the lungs^53^. OMVs derived from *Salmonella typhimurium* and *Escherichia coli* have recently also been used to present respectively the spike RBD or peptides derived from it, and in both cases immunization provided a protective response in animal models^54,55^. In our OMV-mC-Spike vaccine the full length spike protein is included. It has been shown previously that vaccination with S1 induced higher IgG and IgA antibody levels than RBD alone and these antibodies more efficiently neutralized SARS-CoV-2, suggesting many neutralizing epitopes are located in the S1 region and not only in RBD^56^. Another study compared several spike variants, including RBD, S1, S2 and full length spike, and showed that full length spike generated a higher neutralizing-to-binding ratio indicating immunization with the full protein induced more neutralizing antibodies^57^.

A significant limitation for the use of viral vector-based vaccines is the development of anti-vector immunity, which restricts the number of possible booster vaccinations. With OMV based vaccines we assume this is less of an issue, as they are non-replicating and do not depend on a specific host receptor for cellular entry. In addition, major antigens can be removed by gene deletion, as we did with the immunodominant PorA outer membrane protein in our vaccine strain, thus limiting the response against neisserial antigens. OMV-spike vaccination might thus also find an application as a heterologous boost after primary vaccination with viral vector-based vaccines.

Overall, here we show that OMV-mC-Spike is safe and effective in both mice and hamsters. In these animal models, intranasal vaccination with OMV-mC-Spike is superior to intramuscular vaccination, since the amount of IgG induced is higher and in addition a strong mucosal response is induced. Our study demonstrates that adding an mCRAMP tag to the Spike protein and combining it with meningococcal OMVs improves its immunogenicity, thus warranting further development of this vaccine concept towards clinical trials in humans.

## ACKNOWLEDGEMENTS

This work was funded by the Dutch Ministry of Health, Welfare and Sport. The funding source had no role in the design of the study, the analysis and interpretation of the data or the writing of the manuscript. This work was supported by many people at Intravacc, including the OMV production team, the product characterization team and the immunology team.

## AUTHOR CONTRIBUTIONS

ER, DO, and CK. designed and directed the project. AZ designed and performed the experiments and analyzed the data. PL conceptually designed the use of the mCRAMP tag for OMV association, and drafted the first version of the manuscript. All authors provided critical feedback and helped shape the research, analysis and manuscript.

## COMPETING INTERESTS

All authors are employees of Intravacc B.V. The authors attest that the work contained in this research report is free of any bias that might be associated with the commercial goals of the company. Intravacc B.V. filed an international patent application on 07 May 2021: Click OMVs which has received the following application number, PCT/EP2021/062092. PL, AZ, and CK are listed as co-inventors.

