## Supplementary figures for "An intranasal OMV-based vaccine induces high mucosal and systemic protecting immunity against a SARS-CoV-2 infection"

**Supplementary material**

```
Hexapro         MFVFLVLLPLVSSQCVNLTTRTQLPPAYTNSFTRGVYYPDKVFRSSVLHSTQDLFLPFFS   60
Spike           MFVFLVLLPLVSSQCVNLTTRTQLPPAYTNSFTRGVYYPDKVFRSSVLHSTQDLFLPFFS   60
mC-Spike        MFVFLVLLPLVSSQCVNLTTRTQLPPAYTNSFTRGVYYPDKVFRSSVLHSTQDLFLPFFS   60
                ************************************************************

Hexapro         NVTWFHAIHVSGTNGTKRFDNPVLPFNDGVYFASTEKSNIIRGWIFGTTLDSKTQSLLIV   120
Spike           NVTWFHAIHVSGTNGTKRFDNPVLPFNDGVYFASTEKSNIIRGWIFGTTLDSKTQSLLIV   120
mC-Spike        NVTWFHAIHVSGTNGTKRFDNPVLPFNDGVYFASTEKSNIIRGWIFGTTLDSKTQSLLIV   120
                ************************************************************

Hexapro         NNATNVVIKVCEFQFCNDPFLGVYYHKNNKSWMESEFRVYSSANNCTFEYVSQPFLMDLE   180
Spike           NNATNVVIKVCEFQFCNDPFLGVYYHKNNKSWMESEFRVYSSANNCTFEYVSQPFLMDLE   180
mC-Spike        NNATNVVIKVCEFQFCNDPFLGVYYHKNNKSWMESEFRVYSSANNCTFEYVSQPFLMDLE   180
                ************************************************************

Hexapro         GKQGNFKNLREFVFKNIDGYFKIYSKHTPINLVRDLPQGFSALEPLVDLPIGINITRFQT   240
Spike           GKQGNFKNLREFVFKNIDGYFKIYSKHTPINLVRDLPQGFSALEPLVDLPIGINITRFQT   240
mC-Spike        GKQGNFKNLREFVFKNIDGYFKIYSKHTPINLVRDLPQGFSALEPLVDLPIGINITRFQT   240
                ************************************************************

Hexapro         LLALHRSYLTPGDSSSGWTAGAAAYYVGYLQPRTFLLKYNENGTITDAVDCALDPLSETK   300
Spike           LLALHRSYLTPGDSSSGWTAGAAAYYVGYLQPRTFLLKYNENGTITDAVDCALDPLSETK   300
mC-Spike        LLALHRSYLTPGDSSSGWTAGAAAYYVGYLQPRTFLLKYNENGTITDAVDCALDPLSETK   300
                ************************************************************

Hexapro         CTLKSFTVEKGIYQTSNFRVQPTESIVRFPNITNLCPFGEVFNATRFASVYAWNRKRISN   360
Spike           CTLKSFTVEKGIYQTSNFRVQPTESIVRFPNITNLCPFGEVFNATRFASVYAWNRKRISN   360
mC-Spike        CTLKSFTVEKGIYQTSNFRVQPTESIVRFPNITNLCPFGEVFNATRFASVYAWNRKRISN   360
                ************************************************************

Hexapro         CVADYSVLYNSASFSTFKCYGVSPTKLNDLCFTNVYADSFVIRGDEVRQIAPGQTGKIAD   420
Spike           CVADYSVLYNSASFSTFKCYGVSPTKLNDLCFTNVYADSFVIRGDEVRQIAPGQTGKIAD   420
mC-Spike        CVADYSVLYNSASFSTFKCYGVSPTKLNDLCFTNVYADSFVIRGDEVRQIAPGQTGKIAD   420
                ************************************************************

Hexapro         YNYKLPDDFTGCVIAWNSNNLDSKVGGNYNYLYRLFRKSNLKPFERDISTEIYQAGSTPC   480
Spike           YNYKLPDDFTGCVIAWNSNNLDSKVGGNYNYLYRLFRKSNLKPFERDISTEIYQAGSTPC   480
mC-Spike        YNYKLPDDFTGCVIAWNSNNLDSKVGGNYNYLYRLFRKSNLKPFERDISTEIYQAGSTPC   480
                ************************************************************

Hexapro         NGVEGFNCYFPLQSYGFQPTNGVGYQPYRVVVLSFELLHAPATVCGPKKSTNLVKNKCVN   540
Spike           NGVEGFNCYFPLQSYGFQPTNGVGYQPYRVVVLSFELLHAPATVCGPKKSTNLVKNKCVN   540
mC-Spike        NGVEGFNCYFPLQSYGFQPTNGVGYQPYRVVVLSFELLHAPATVCGPKKSTNLVKNKCVN   540
                ************************************************************

Hexapro         FNFNGLTGTGVLTESNKKFLPFQQFGRDIADTTDAVRDPQTLEILDITPCSFGGVSVITP   600
Spike           FNFNGLTGTGVLTESNKKFLPFQQFGRDIADTTDAVRDPQTLEILDITPCSFGGVSVITP   600
mC-Spike        FNFNGLTGTGVLTESNKKFLPFQQFGRDIADTTDAVRDPQTLEILDITPCSFGGVSVITP   600
                ************************************************************

Hexapro         GTNTSNQVAVLYQDVNCTEVPVAIHADQLTPTWRVYSTGSNVFQTRAGCLIGAEHVNNSY   660
Spike           GTNTSNQVAVLYQGVNCTEVPVAIHADQLTPTWRVYSTGSNVFQTRAGCLIGAEHVNNSY   660
mC-Spike        GTNTSNQVAVLYQGVNCTEVPVAIHADQLTPTWRVYSTGSNVFQTRAGCLIGAEHVNNSY   660
                *************.**********************************************

Hexapro         ECDIPIGAGICASYQTQTNSPGSASSVASQSIIAYTMSLGAENSVAYSNNSIAIPTNFTI   720
Spike           ECDIPIGAGICASYQTQTNSPGSASSVASQSIIAYTMSLGAENSVAYSNNSIAIPTNFTI   720
mC-Spike        ECDIPIGAGICASYQTQTNSPGSASSVASQSIIAYTMSLGAENSVAYSNNSIAIPTNFTI   720
                ************************************************************

Hexapro         SVTTEILPVSMTKTSVDCTMYICGDSTECSNLLLQYGSFCTQLNRALTGIAVEQDKNTQE   780
Spike           SVTTEILPVSMTKTSVDCTMYICGDSTECSNLLLQYGSFCTQLNRALTGIAVEQDKNTQE   780
mC-Spike        SVTTEILPVSMTKTSVDCTMYICGDSTECSNLLLQYGSFCTQLNRALTGIAVEQDKNTQE   780
                ************************************************************
```

```
Hexapro      VFAQVKQIYKTPPIKDFGGFNFSQILPDPSKPSKRSPIEDLLFNKVTLADAGFIKQYGDC  840
Spike        VFAQVKQIYKTPPIKDFGGFNFSQILPDPSKPSKRSPIEDLLFNKVTLADAGFIKQYGDC  840
mC-Spike     VFAQVKQIYKTPPIKDFGGFNFSQILPDPSKPSKRSPIEDLLFNKVTLADAGFIKQYGDC  840
             ************************************************************

Hexapro      LGDIAARDLICAQKFNGLTVLPPLLTDEMIAQYTSALLAGTITSGWTFGAGPALQIPFPM  900
Spike        LGDIAARDLICAQKFNGLTVLPPLLTDEMIAQYTSALLAGTITSGWTFGAGPALQIPFPM  900
mC-Spike     LGDIAARDLICAQKFNGLTVLPPLLTDEMIAQYTSALLAGTITSGWTFGAGPALQIPFPM  900
             ************************************************************

Hexapro      QMAYRFNGIGVTQNVLYENQKLIANQFNSAIGKIQDSLSSTPSALGKLQDVVNQNAQALN  960
Spike        QMAYRFNGIGVTQNVLYENQKLIANQFNSAIGKIQDSLSSTPSALGKLQDVVNQNAQALN  960
mC-Spike     QMAYRFNGIGVTQNVLYENQKLIANQFNSAIGKIQDSLSSTPSALGKLQDVVNQNAQALN  960
             ************************************************************

Hexapro      TLVKQLSSNFGAISSVLNDILSRLDPPEAEVQIDRLITGRLQSLQTYVTQQLIRAAEIRA  1020
Spike        TLVKQLSSNFGAISSVLNDILSRLDPPEAEVQIDRLITGRLQSLQTYVTQQLIRAAEIRA  1020
mC-Spike     TLVKQLSSNFGAISSVLNDILSRLDPPEAEVQIDRLITGRLQSLQTYVTQQLIRAAEIRA  1020
             ************************************************************

Hexapro      SANLAATKMSECVLGQSKRVDFCGKGYHLMSFPQSAPHGVVFLHVTYVPAQEKNFTTAPA  1080
Spike        SANLAATKMSECVLGQSKRVDFCGKGYHLMSFPQSAPHGVVFLHVTYVPAQEKNFTTAPA  1080
mC-Spike     SANLAATKMSECVLGQSKRVDFCGKGYHLMSFPQSAPHGVVFLHVTYVPAQEKNFTTAPA  1080
             ************************************************************

Hexapro      ICHDGKAHFPREGVFVSNGTHWFVTQRNFYEPQIITTDNTFVSGNCDVVIGIVNNTVYDP  1140
Spike        ICHDGKAHFPREGVFVSNGTHWFVTQRNFYEPQIITTDNTFVSGNCDVVIGIVNNTVYDP  1140
mC-Spike     ICHDGKAHFPREGVFVSNGTHWFVTQRNFYEPQIITTDNTFVSGNCDVVIGIVNNTVYDP  1140
             ************************************************************

Hexapro      LQPELDSFKEELDKYFKNHTSPDVDLGDISGINASVVNIQKEIDRLNEVAKNLNESLIDL  1200
Spike        LQPELDSFKEELDKYFKNHTSPDVDLGDISGINASVVNIQKEIDRLNEVAKNLNESLIDL  1200
mC-Spike     LQPELDSFKEELDKYFKNHTSPDVDLGDISGINASVVNIQKEIDRLNEVAKNLNESLIDL  1200
             ************************************************************

Hexapro      QELGKYEQGSGYIPEAPRDGQAYVRKDGEWVLLSTFLGRSLEVLFQGPGHHHHHHHHSAW  1260
Spike        QELGKYEQGSGYIPEAPRDGQAYVRKDGEWVLLSTFLGRSLEVLFQGPGHHHHHHHHSAW  1260
mC-Spike     QELGKYEQGSGYIPEAPRDGQAYVRKDGEWVLLSTFLGRSLEVLFQGPGHHHHHHHHSAW  1260
             ************************************************************

Hexapro      SHPQFEKGGGSGGGGSGGSAWSHPQFEK---------------------------------  1288
Spike        SHPQFEKGGGSG-GGSGGSAWSHPQFEK--------------------------------  1287
mC-Spike     SHPQFEKGGGSG-GGSGGSAWSHPQFEKGGGSGGGSGGGSGLLRKGGEKIGEKLKKIGQK  1319
             ************ ***************

Hexapro      --------------      1288
Spike        --------------      1287
mC-Spike     IKNFFQKLVPQPEQ      1333
```

**Figure S1. Multiple sequence alignment of the Spike protein sequences.** HexaPro spike protein (HexaPro) expression construct sequence (DOI: 10.2210/pdb6XKL/pdb) was aligned with the sequence of the spike protein (mC-Spike) in the OMV-mC-Spike vaccine and the Spike protein (Spike) that was mixed with OMVs (OMV+Spike) in the mouse study. Both Spike and mC-Spike protein contain the D614G mutation not present in HexaPro. The mC-Spike protein contains a 3XGGGS repeat and a short amphipathic peptide, mCramp (GenBank accession number: CAA64078), an LPS-binding tag which are lacking in the Spike protein and HexaPro.

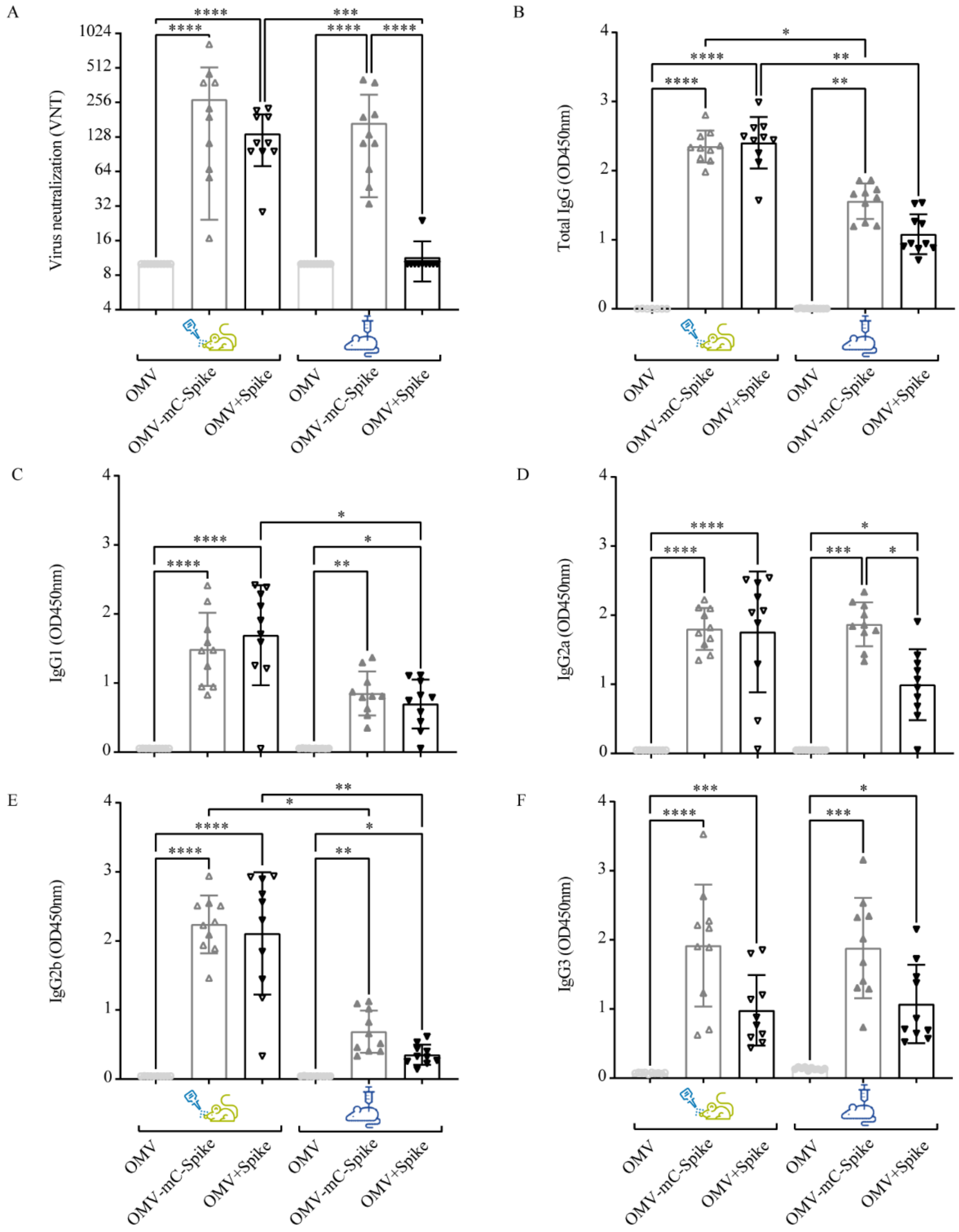


**Figure S2. Induction of virus neutralizing antibody titers and spike IgG antibody responses in mice.** Balb/C mice were immunized intranasally or intramuscularly on day 0 and day 21 with 15µg OMV (control group) or 15 µg OMV combined with 15µg Spike with mCRAMP (OMV-mC-Spike) or without mCRAMP (OMV+Spike). Sera was collected at day 35 and virus neutralizing titers (**A**) were determined. Furthermore spike IgG antibody levels were measured at day 35 with ELISA. Total IgG antibody levels were determined in sera diluted at 1:50000 (**B**). IgG subtypes IgG1 (**C**), IgG2a (**D**), IgG2b (**E**) and IgG3 (**F**) were measured in sera respectively diluted at 1:50000 (IgG1), 1:10000 (IgG2a), 1:50000 (IgG2b) and 1:100 (IgG3). Statistical significance was determined using the Kruskal-Wallis test followed by Two-stage linear step-up procedure of Benjamini, Krieger and Yekutieli multiple-comparison test. Significance is depicted as *p<0.05, **p<0.01***p < 0.001, **** p<0.0001.

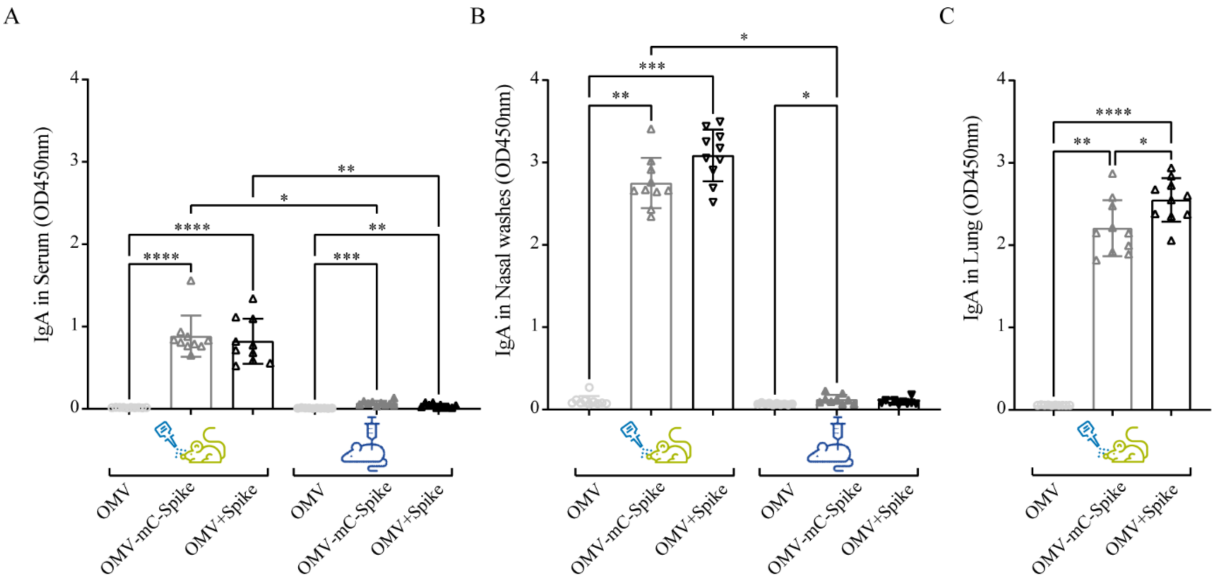


**Figure S3. IgA antibody responses after intranasal immunization in mice.** Balb/C mice were immunized intranasally or intramuscularly on day 0 and day 21 with 15µg OMV (control group) or 15 mg OMV combined with 15 mg Spike with mCRAMP (OMV-mC-Spike) or without mCRAMP (OMV+Spike). At day 35 sera and nasal washes were collected from all animals, and lungs were collected from animals which were intranasally immunized. IgA antibodies were measured with an ELISA in serum (1:200 dilution) (**A** ), nasal wash (1:1 dilution) (**B**) and lung (1:50 dilution) (**C**). Statistical significance was determined using the Kruskal-Wallis test followed by a two-stage linear step-up procedure of Benjamini, Krieger and Yekutieli multiple-comparison test. Significance is depicted as *p<0.05, **p<0.01***p < 0.001, **** p<0.0001.

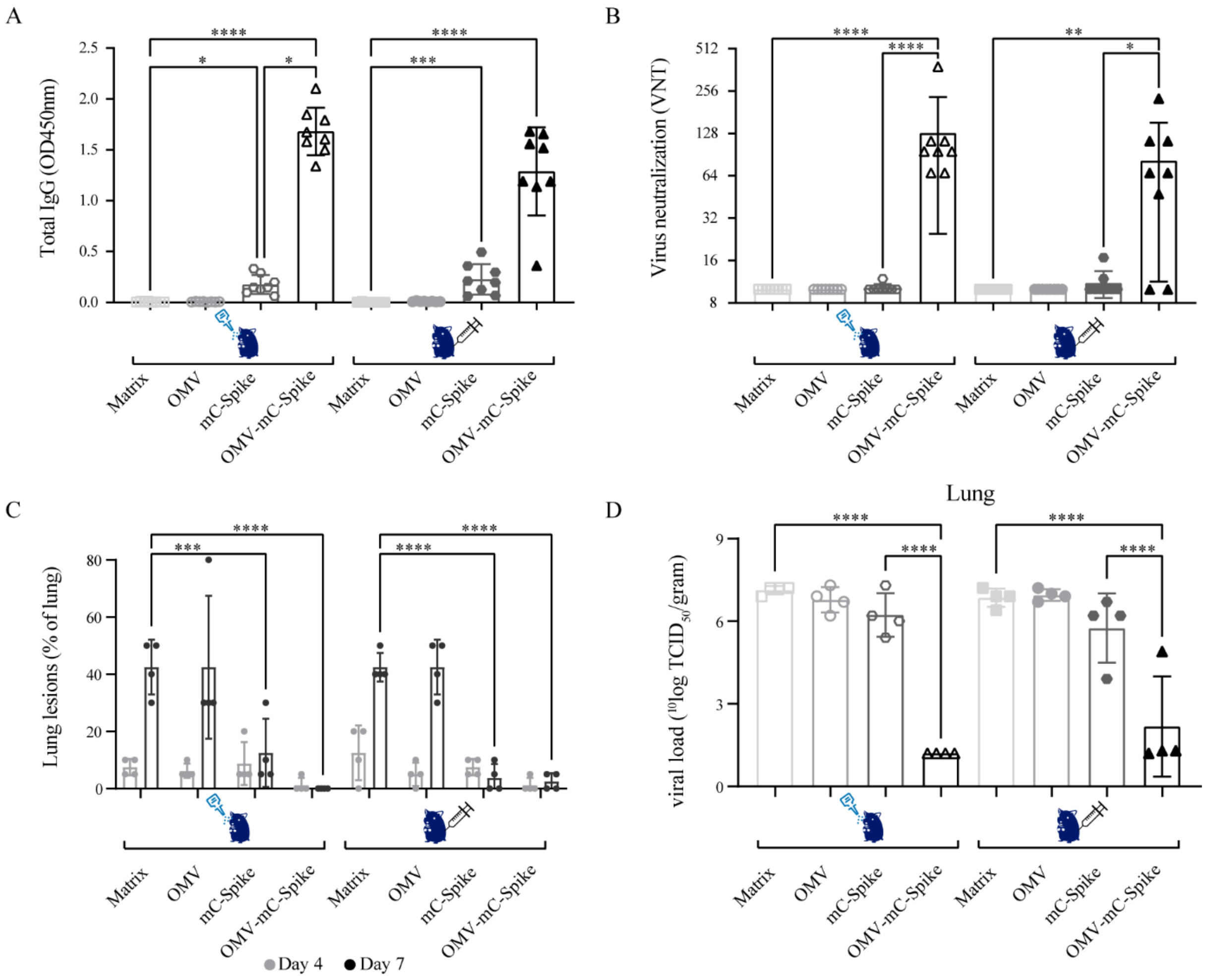


**Figure S4. Vaccination with OMV-mC-Spike protects against SARS-CoV-2 in Syrian hamsters.** Animals were immunized intranasally or intramuscularly on day 0 and day 21 with 15µg OMV, 15µg spike with mCRAMP (mC-Spike) or 15µg OMV combined with 15µg mC-Spike (OMC-mC-Spike). In the control group animals were immunized with 10mM Tris-3% sucrose, which is the OMV buffer. Sera were collected from all hamsters at experimental day 0, 21, 42, 46 and 49. At day 46 half of the animals per group (4 out of 8) were sacrificed and at day 49 the remaining 4 animals were sacrificed. (**A**) From the sera at day 42 total spike IgG antibodies were determined with an ELISA. Sera was 1:4000 diluted. (**B**) Virus neutralizing titers were determined in sera from day 42. (**C**) When animals were sacrificed at day 46 (day 4 post challenge) and day 49 (Day 7 post challenge) the percentage of the lung that presented lung lesions was quantified. The viral load was determined in lungs (**D**), throat swabs and nasal turbinates (not shown). Statistical significance of the difference was evaluated by Kruskal-Wallis test and followed by a two-stage linear step-up procedure of Benjamini, Krieger and Yekutieli multiple-comparison test (**A, B** and **D**) or by a 2-way ANOVA test followed by Tukey's multiple-comparison test (**C**). Significance is depicted as *p<0.05, **p<0.01***p < 0.001, **** p<0.0001.

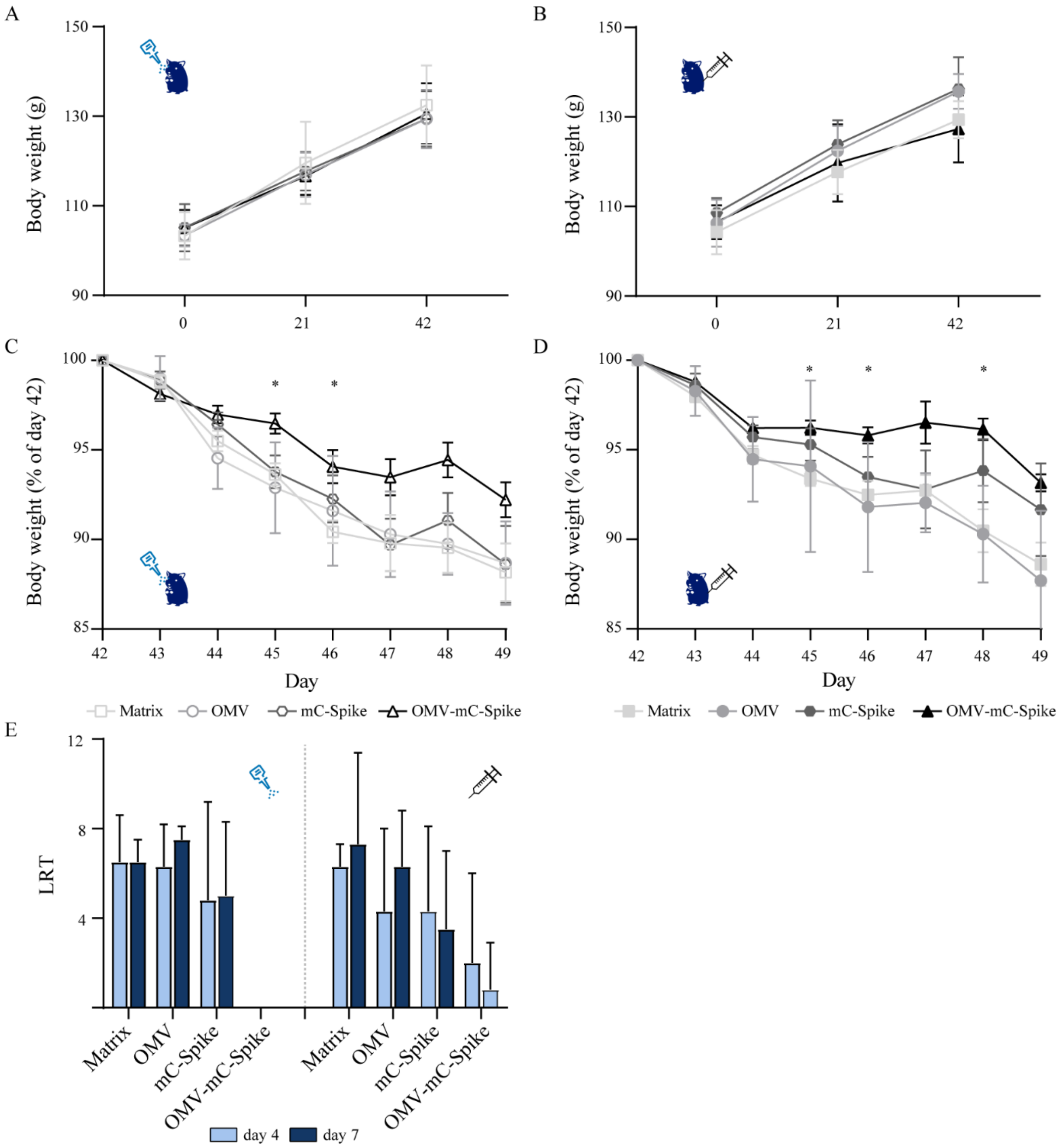


**Figure S5. Body weight and histopathological analysis.** Animals were immunized as described in figure S4. Bodyweight was measured at day 0, 21 and 42 in intranasal (**A**) and intramuscular (**B**) immunized animals prior to challenge with SARS-CoV-2. After challenge, the bodyweight was daily measured in intranasal (**C**) and intramuscular (**D**) immunized animals. Day 47-49 only include 4 animals, because half of the animals were sacrificed at day 46. Histopathological analysis from lung tissues was performed for all animals (**E**). Lung tissue was analyzed and scored for presence and severity of alveolitis, alveolar damage, alveolar edema, alveolar hemorrhage, type II pneumocyte hyperplasia, bronchitis, bronchiolitis, peribronchial and perivascular cuffing. Scores are presented as sum of Lower Respiratory Tract (LRT) disease parameters. Statistical significance was determined with a 2-way ANOVA test followed by Tukey's multiple-comparison. Significance is depicted as *p<0.05, **p<0.01***p < 0.001, **** p<0.0001.
